# Mlp1 and Mlp2 cooperate to build a stoichiometric nuclear pore basket in budding yeast

**DOI:** 10.64898/2026.03.13.709474

**Authors:** Keri Schmidt, Alexandra P Schürch, Elisa Dultz

**Affiliations:** Institute of Biochemistry, Department of Biology, ETH Zurich, Zurich 8092, Switzerland; Departments of Biology and Chemistry, Utrecht University, 3584 CG Utrecht, the Netherlands

**Keywords:** Nuclear pore complex, nuclear pore basket, S. cerevisiae, Mlp1, Mlp2

## Abstract

The nuclear pore complex is the only gateway between the nucleus and the cytoplasm in eukaryotic cells. Its nucleoplasmic face is decorated by the nuclear basket, a filamentous structure with important roles in mRNA export and chromatin organization. In contrast to major parts of the nuclear pore scaffold, the architecture of the nuclear basket remains poorly defined. Here, we investigate the interactions required for formation and maintenance of the nuclear basket *in vivo* using budding yeast. While previous work has often focused on Mlp1, the largest and most abundant nuclear basket protein, we demonstrate that its paralogue, Mlp2, also plays a central role in nuclear basket architecture. Mlp2 can interact with the NPC scaffold independently of Mlp1, and interactions between the coiled-coil regions of both proteins stabilize their binding. Furthermore, the N-termini of both Mlp1 and Mlp2 are required for recruitment of Pml39. In addition, we show that Pml39 uses its N- and C-terminal helices to recruit additional Mlp1 subunits. We propose a refined model of nuclear basket architecture with a stoichiometry of 4:2:1 per spoke for Mlp1:Mlp2:Pml39.

**Summary statement:** Based on experiments in budding yeast, a refined model for nuclear pore basket architecture is proposed with a stoichiometry of 4:2:1 per spoke for the major components Mlp1:Mlp2:Pml39.

## Introduction

The nuclear pore complex (NPC) is a massive, proteinaceous channel embedded in the nuclear envelope that serves as the sole transport passage between the nucleus and the cytoplasm. Composed of a family of proteins known as nucleoporins (Nups), the NPC core consists of a stack of three eight-fold symmetric rings that form the central channel (Fig. 1A). The outer rings are decorated with asymmetric, filamentous features that have compartment-specific functions: At the cytoplasmic face, the mRNA export platform remodels mRNPs as they exit the channel, while at the nucleoplasmic face, a mesh-like structure known as the nuclear basket plays critical roles in mRNA export as well as non-transport related functions, including chromatin organization and genome maintenance (reviewed in (De Magistris, 2021; Dultz and Doye, 2025; Nobari et al., 2023; Simon et al., 2024).

**Figure 1:**
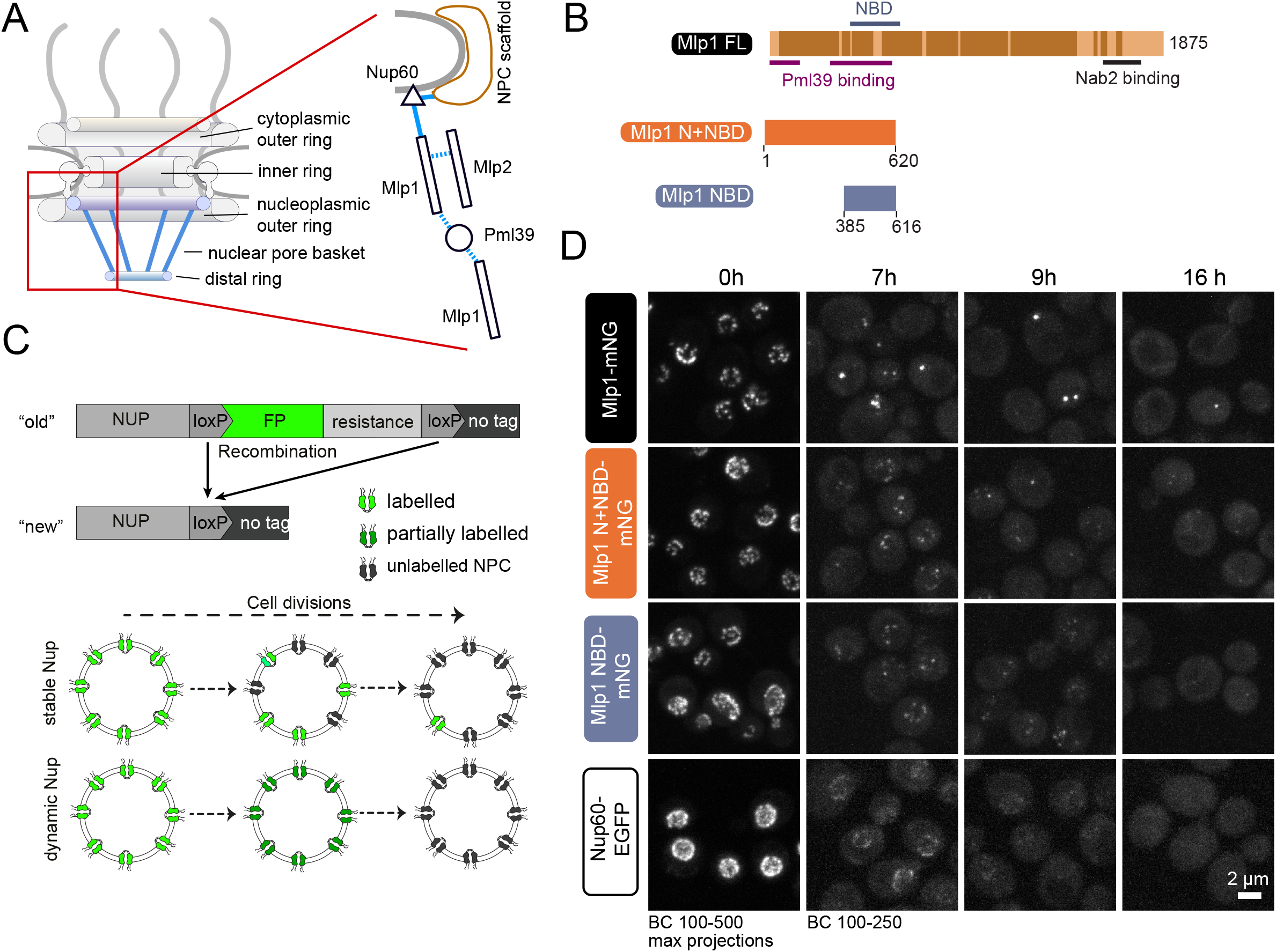
Mlp1 truncations bind the NPC stably. (A)Schematic representation of the nuclear pore complex and in more detail the nuclear basket composition. Dashed blue lines indicate characterized direct interactions. The solid lines connecting Nup60 to the NPC scaffold and Mlp1 indicate that these interactions are structurally characterized at the residue level. Left part of figure was adapted from (Dultz and Doye, 2025). (B)Schematic representation of full-length and truncated Mlp1 constructs used in this study. Darker shaded areas on the full-length protein representing Mlp1 represent predicted coiled-coil domains of the protein (Gabler et al., 2020). Characterized interaction regions with Pml39, Nup60 and Nab2 are indicated. (C)Schematic representation of doRITE: A nucleoporin of interest is tagged with a fluorescent protein (FP). Upon induction of Cre-recombinase, newly synthesized NUP will no longer be fluorescent. As cells divide, this leads to the dilution of labelled NPCs for stable Nups, while dynamic Nups redistribute between new and old NPCs and thus a decrease of the signal at the nuclear envelope is observed. Adapted from (Zsok et al., 2024). (D)Representative image panels from a doRITE assay carried out with cells expressing the different Mlp1 truncation constructs introduced in (B) tagged with the RITE cassette at different timepoints after RITE induction. All Mlp1 constructs shown carry an N-terminal nuclear localization signal (NLS). Note that the contrast settings for image panels of the 0 h timepoint are different from the contrast settings of the remaining panels. The image panels shown in this figure are maximum intensity projections.

While recent structural studies have resolved the majority of the NPC scaffold and the cytoplasmic export platform to high resolution (reviewed in (Huang et al., 2023; Petrovic et al., 2022)), the nuclear basket remains structurally elusive. Its high proportion of intrinsically disordered domains and apparent inherent flexibility have, to date, hampered high-resolution analysis, leaving its precise architecture unclear.

In yeast, the primary structural components of the nuclear basket are the paralogues Mlp1 and Mlp2 (TPR in mammals) (Fig. 1A shows an overview of the nuclear pore basket components and their interactions). These large, coiled-coil filamentous proteins anchor to the NPC via a central domain with both termini extending into the nucleoplasm (Krull et al., 2004; Niepel et al., 2013; Strambio-de-Castillia et al., 1999). *In vitro*, Mlp1 dimers have been visualized as extended rods with flexible joints, but extensive internal cross-links in the protein suggest that it might fold back and forth on itself (Kim et al., 2018; Stankunas and Köhler, 2024). Cross-links also suggest that Mlp1 and Mlp2 are arranged in a parallel manner (Kim et al., 2018), with Mlp2 depending on Mlp1 for recruitment to the NPC (Palancade et al., 2005). Recent cryo-electron tomography has identified density for these filaments and pinpointed their anchor sites at the nuclear outer ring (Akey et al., 2022; Singh et al., 2024), but the resolution remains insufficient to precisely define the interactions with the scaffold and to determine how the coiled-coil regions of Mlp1 and Mlp2 are organized individually and relative to one another.

During assembly of the NPC in budding yeast, association of the nuclear pore basket filaments is a very late step (Onischenko et al., 2020). The initial recruitment of the filaments is mediated by the disordered nucleoporin Nup60 (NUP153 in mammals), which utilizes multiple short sequence motifs to bridge Mlp1 to the NPC scaffold and the nuclear envelope (Cibulka et al., 2022; Mészáros et al., 2015; Stankunas and Köhler, 2024). However, additional uncharacterized interactions between the filaments and the NPC must exist, given that Nup60 becomes dispensable for their maintenance at the NPC once the basket is assembled (Aksenova et al., 2020; King et al., 2023; Zsok et al., 2024). Further copies of Mlp1 are recruited to the nuclear basket by Pml39 (ZC1HC3 in mammals), which was only recently recognized as a *bona fide* Nup that can bind both Mlp2 and two distinct regions on Mlp1 (Gunkel et al., 2023, 2021; Gunkel and Cordes, 2022; Palancade et al., 2005) (Fig. 1B). How Pml39 orchestrates this conserved interlinkage and contributes to basket architecture remains unclear.

The nuclear basket acts as a platform for the recruitment of several specific factors including the poly A binding protein Nab2 and the TREX-2 complex, both involved in mRNA export, or the Sumo-protease Ulp1, which is responsible for the bulk of de-sumoylation reactions in budding yeast (McNeil et al., 2024). Intriguingly, in budding yeast only a subset of NPCs carry a nuclear basket (Bensidoun et al., 2022; Galy et al., 2004), thus generating different NPC subpopulations that might perform specialized functions. However, the lack of a detailed architectural model for the nuclear basket has so far prevented a molecular understanding of how it coordinates mRNA export, chromatin organization and gene expression. In this study, we utilized truncation analyses and quantitative microscopy to map the interaction network required for a stable nuclear basket *in vivo* using *S. cerevisiae*. Based on our findings, we propose a refined model for the architecture and assembly of the nuclear basket.

## Results

### The NPC binding domain of Mlp1 binds stably to NPCs over multiple cell divisions

In the budding yeast *S. cerevisiae*, nuclear basket assembly is initiated by the recruitment of Mlp1 through the interaction of its central NPC-binding domain (amino acids (aa) 385-616, Mlp1-NBD) with Nup60 ((Cibulka et al., 2022; Niepel et al., 2013; Stankunas and Köhler, 2024), Fig. 1A,B). Given that Mlp1 remains stably bound to the NPC over multiple cell divisions (Onischenko et al., 2020; Hakhverdyan et al., 2021; Zsok et al., 2024), we wanted to determine whether the Mlp1-NBD is sufficient for mediating this long-term, stable interaction. To examine the stability of Mlp1 truncation mutants at the NPC, we employed the doRITE assay (dilution of ‹recombination induced tag exchange›-labelled complexes), which distinguishes between stable and dynamic Nup associations (Zsok et al., 2024) (Fig. 1C). In doRITE, Mlp1 constructs are initially expressed with a fluorescent tag. Upon activation of Cre recombinase, the fluorescent tag is excised from the genome and all newly synthesized proteins are thus non-fluorescent, meaning that “old” protein can be followed exclusively. Nups that are stably associated with individual NPCs exhibit a characteristic punctate pattern where the number of dots is reduced in each cell division as “old”, labeled NPCs are partitioned between mother and daughter cells. In contrast, dynamic Nups continuously redistribute between different NPCs, resulting in a signal that becomes uniformly dimmer across the entire NPC population (Fig. 1C).

Our doRITE analysis revealed that the Mlp1-NBD alone or a construct encompassing in addition the N-terminus (N+NBD) exhibit stable binding over multiple generations, as evidenced by the dilution of labelled NPC foci with consecutive cell divisions (Figs 1D, S1A). Defined albeit dim NPC foci could be observed even after 16 h, representing more than seven cell divisions. This is in contrast to the dynamic Nup60, which exhibited no punctate staining and no signal over background could be observed after 16 h. Thus, the NBD (aa385-616) is sufficient for stable association of Mlp1 with the NPC and the coiled-coil segments that likely form the basket filaments are dispensable.

Although the signal intensity of individual NPC foci was reduced in cells expressing the truncation constructs when compared to the full-length protein, the foci number was higher at all time points (Fig. S1A). Of note, the truncated constructs were expressed at higher levels than the wild-type copy (Fig. S1B). Therefore, the increased number of foci is likely a consequence of higher expression, which leads to a larger soluble pool from which fluorescent subunits can be integrated into newly synthesized NPCs for an extended period after RITE induction.

### The C-terminus of Mlp1 confers stable binding in the absence of Nup60

Nup60 is important for the initial recruitment of Mlp1, and its deletion typically results in Mlp1 mislocalization to nucleoplasmic foci (Feuerbach et al., 2002; Galy et al., 2004). However, it was recently shown that localization of Mlp1 to NPCs in *nup60Δ* cells is partially restored when cells are grown at lower temperature (Ptak et al., 2025). Furthermore, once Mlp1 is assembled into the basket, Nup60 becomes dispensable for its maintenance (King et al., 2023; Zsok et al., 2024). These observations suggest that Mlp1 has Nup60-independent interactions with the NPC scaffold (Fig. 2A), which we sought to characterize.

**Figure 2:**
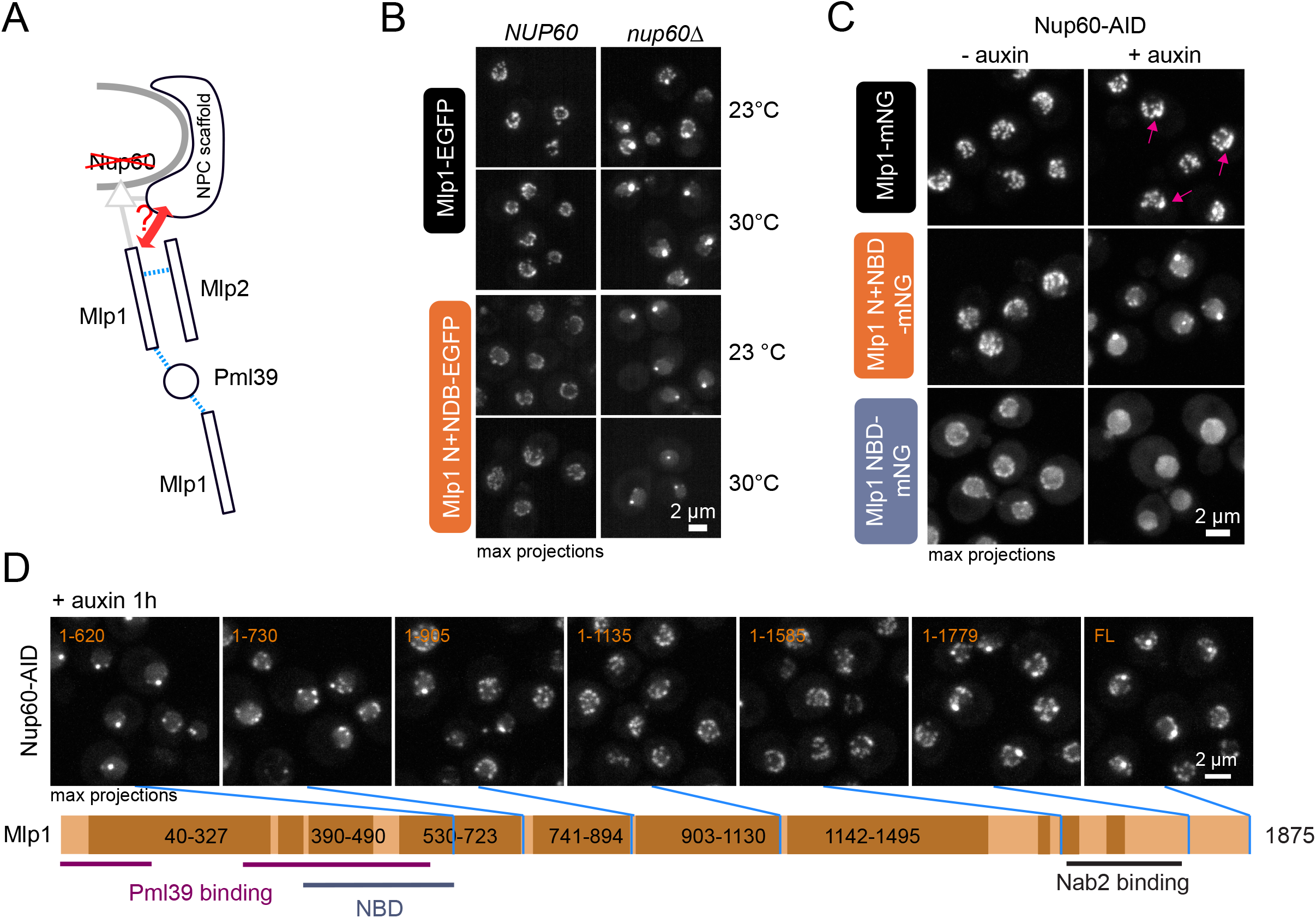
The Mlp1 C-terminus confers Nup60-independent binding at the NPC. (A)Schematic illustration of the interaction analyzed in this figure: Nup60-independent interactions between Mlp1 and the NPC scaffold. (B)Localization of full-length or truncated Mlp1 in the presence or absence of Nup60 at different temperatures. (C)Localization of the different Mlp1 constructs upon acute depletion of Nup60 via an auxin-inducible degron. Auxin treatment was performed for 1h. Pink arrows indicate foci of Mlp1-monomericNeonGreen (mNG). (D)Localization of EGFP tagged, C-terminally truncated versions of Mlp1 after auxin-inducible degradation of Nup60 1 h after auxin addition. Orange numbers describe the amino acids of Mlp1 present in the constructs. Blue lines schematically indicate the truncation position on the full-length protein. The darker orange regions on the Mlp1 schematic represent regions that are predicted to form coiled-coil structures. The image panels shown are maximum intensity projections.

First, we examined the localization of the N+NBD fragment in *nup60Δ* cells at a reduced temperature. As reported, full-length Mlp1 tagged with enhanced green fluorescent protein (EGFP) exhibited more punctate NPC localization when the cells were grown at 23°C instead of 30°C (Fig. 2B). In contrast, the Mlp1 N+NBD fragment mislocalized to the nucleoplasm and formed distinct foci in the absence of Nup60 at both temperatures (Fig. 2B). Thus, a region in the C-terminus of Mlp1 is required for Nup60-independent nuclear basket assembly.

To test whether this also applies for the maintenance of the basket after assembly, we used an auxin-inducible degron (AID) to acutely deplete Nup60 (Fig. S2A)(Nishimura et al., 2009). Upon depletion, full-length Mlp1 remained at the nuclear periphery (Fig. 2C), consistent with prior findings (Zsok et al., 2024). Note that the Mlp1-EGFP foci that form in these conditions consist of newly translated protein that cannot incorporate into NPCs in the absence of Nup60 (Zsok et al., 2024). In contrast, the Mlp1 fragments NBD and N+NBD dissociated from the nuclear periphery upon Nup60 depletion (Fig. 2C). Thus, the interaction of the Mlp1 NBD and N-terminus with the NPC is strictly dependent on Nup60 for both incorporation and stable maintenance. Whereas the NBD was largely diffuse in the nucleus, the N+NBD fragment formed a bright focus. This is consistent with the presence of the N-terminal Pml39 binding site in the N+NBD construct, which promotes interlinking of Mlp1 dimers into larger assemblies (Gunkel et al., 2023).

To identify the specific C-terminal regions that mediate the Nup60-independent NPC interaction, we analyzed a series of C-terminally truncated Mlp1 variants following acute Nup60 depletion. We observed a length-dependent improvement in NPC association, which did not correlate with the expression level of the constructs (Figs 2D, S2B). While the N+NBD fragment showed only residual peripheral localization one hour after induction of Nup60 depletion, fragments extending to aa 730 and aa 905 showed more NPC-like foci. The fragments aa1-1135 and aa1-1585 exhibited localizations comparable to the undepleted controls. Interestingly, the longest constructs (aa 1–1779 and full-length) retained NPC localization but formed additional bright nucleoplasmic foci, possibly because of the C-terminal Nab2 (nuclear poly A binding protein) binding site (Fasken et al., 2008), which might promote focus formation via mRNP-mediated interactions. However, other features of the disordered C-terminus or differences in expression levels could also contribute (Fig. S2B). A similar stabilization of NPC association was observed with increasing fragment lengths in cells grown in conditions of constant Nup60 depletion at reduced temperature (Fig. S2C). Together these findings suggest that the C-terminal coiled-coil region of Mlp1 contains multiple segments important for Nup60-independent NPC interaction.

### Mlp1 and Mlp2 mutually stabilize each other at the NPC

Cross-linking mass-spectrometry suggests that Mlp2, the less-characterized paralogue of Mlp1, binds to the basket in a parallel orientation to Mlp1 (Kim et al., 2018). We therefore hypothesized that lateral interactions between the paralogues might stabilize the basket filaments upon depletion of Nup60 (Fig. 3A). Mlp2 shares a similar organization with Mlp1, but it was thought to rely on Mlp1 for localization to the NPC (Palancade et al., 2005). However, we found that Mlp2 also localizes to the NPC in the absence of Mlp1 (Fig. 3B). A region homologous to the NBD characterized for Mlp1 (Mlp2 aa 354-600) was sufficient and necessary to mediate NPC targeting in the absence of Mlp1. Some residual interaction with NPCs was observed for the construct lacking the NBD if Mlp1 was present (Fig. 3B) but not in the absence of Mlp1.

**Figure 3:**
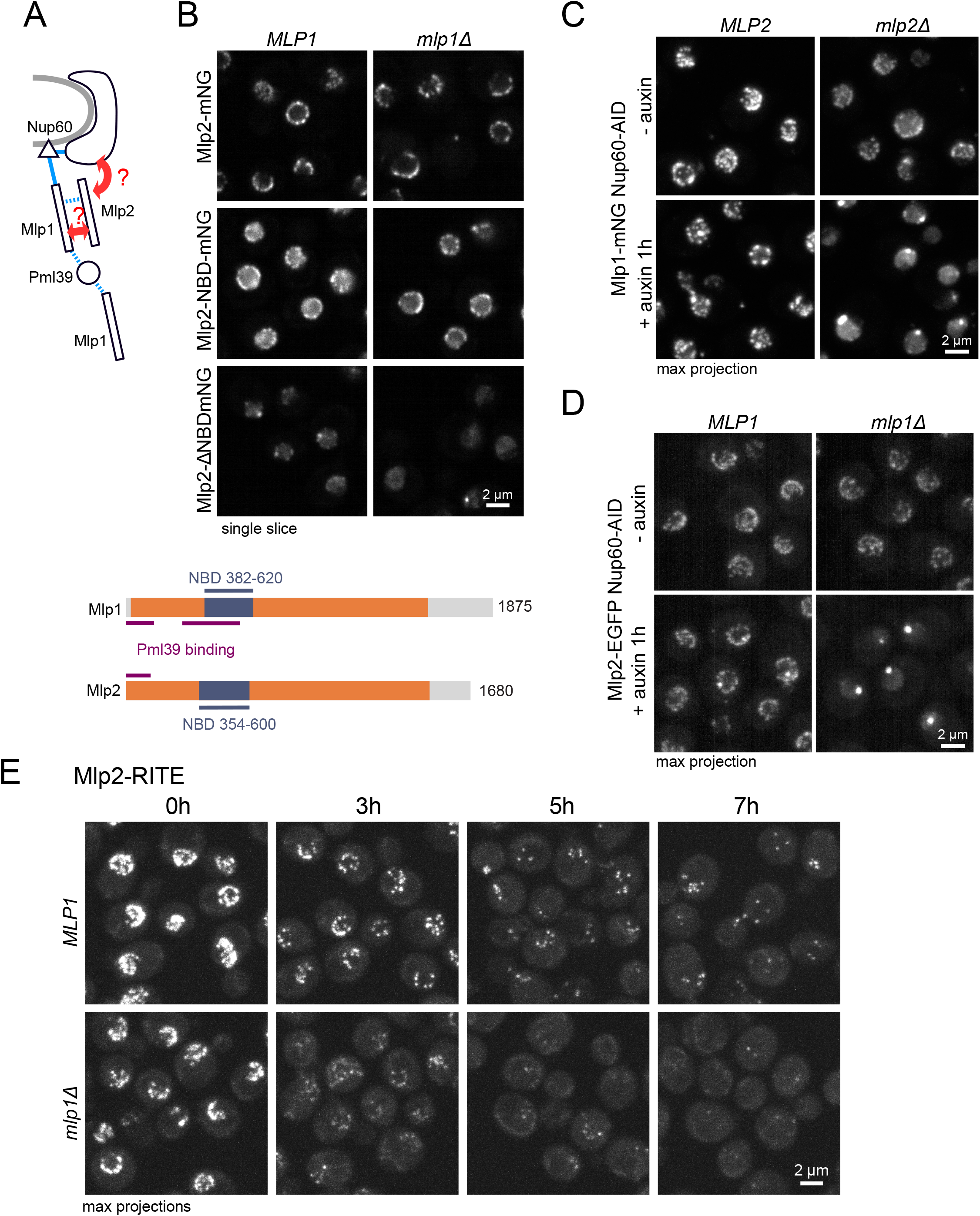
Mlp2 can interact with the NPC on its own but is stabilized there by Mlp1. (A)Schematic illustration of the interaction analyzed in this figure: interaction of Mlp2 with the NPC scaffold and Mlp1. (B)Localization of full-length and truncated constructs of Mlp2 in the presence or absence of Mlp1. Schematic shows homology of Mlp1 and Mlp2. Orange regions align by BlastP (Camacho et al., 2009). Blue regions indicate the NBD of Mlp1 and the corresponding aligned region in Mlp2. (C)Localization of Mlp1 upon depletion of Nup60 via auxin-induced degradation 1h after auxin treatment in the presence or absence of Mlp2. (D)Localization of Mlp2 upon depletion of Nup60 via auxin-induced degradation 1h after auxin treatment in the presence or absence of Mlp1. (E)Representative maximum intensity projections from different time points of a doRITE assay with cells expressing Mlp2 tagged with the RITE cassette in the presence of absence of Mlp1.

Next, we tested the behavior of Mlp1 upon Nup60 depletion when Mlp2 was deleted. Mlp1 readily dissociated from the nuclear basket upon depletion of Nup60 when Mlp2 was absent (Fig. 3C). This could be rescued by expression of full-length Mlp2 but not the NBD, N+NBD or ΔNBD constructs (Fig. S3A). Vice versa, Mlp2 was stable at the NPC upon depletion of Nup60 only in the presence of Mlp1 but readily dissociated in a mlp1Δ background (Fig. 3D). Again, this could be rescued only by a full length Mlp1 but not by the NBD or N+NBD constructs (Fig. S3B). These results suggest that Nup60 supports the interaction of both Mlp1 and Mlp2 at the NPC and that they mutually stabilize each other via interactions of their C-terminal coiled-coil segments.

Using the RITE assay, we next tested whether Mlp1 contributes to long-term stability of Mlp2at the NPC. We found that Mlp2 exhibited strongly reduced stability at the NPC in the absence of Mlp1 as indicated by a drastic drop in signal intensity at NPCs three hours after RITE induction as well as a reduction in number and intensity of NPC foci at later timepoints (Fig. 3E). Thus, Mlp2 can – supported by Nup60 – bind to the NPC via its NBD, but this interaction is not stable in the absence of Mlp1.

### Stoichiometric Mlp1 recruitment to the pore depends on Mlp2 and Pml39

In contrast to Mlp2, Mlp1 remained stably bound to NPC foci in the absence of its paralog (Figs 4A, S4A, S1A). However, we observed a significant decrease in Mlp1-EGFP signal individual NPC foci (Fig. 4A), suggesting that Mlp2 is required to stabilize or recruit a specific subpopulation of Mlp1 at the pore. Deletion of *MLP2* did not affect the total levels of Mlp1 in the cells (Fig. S1B).

**Figure 4:**
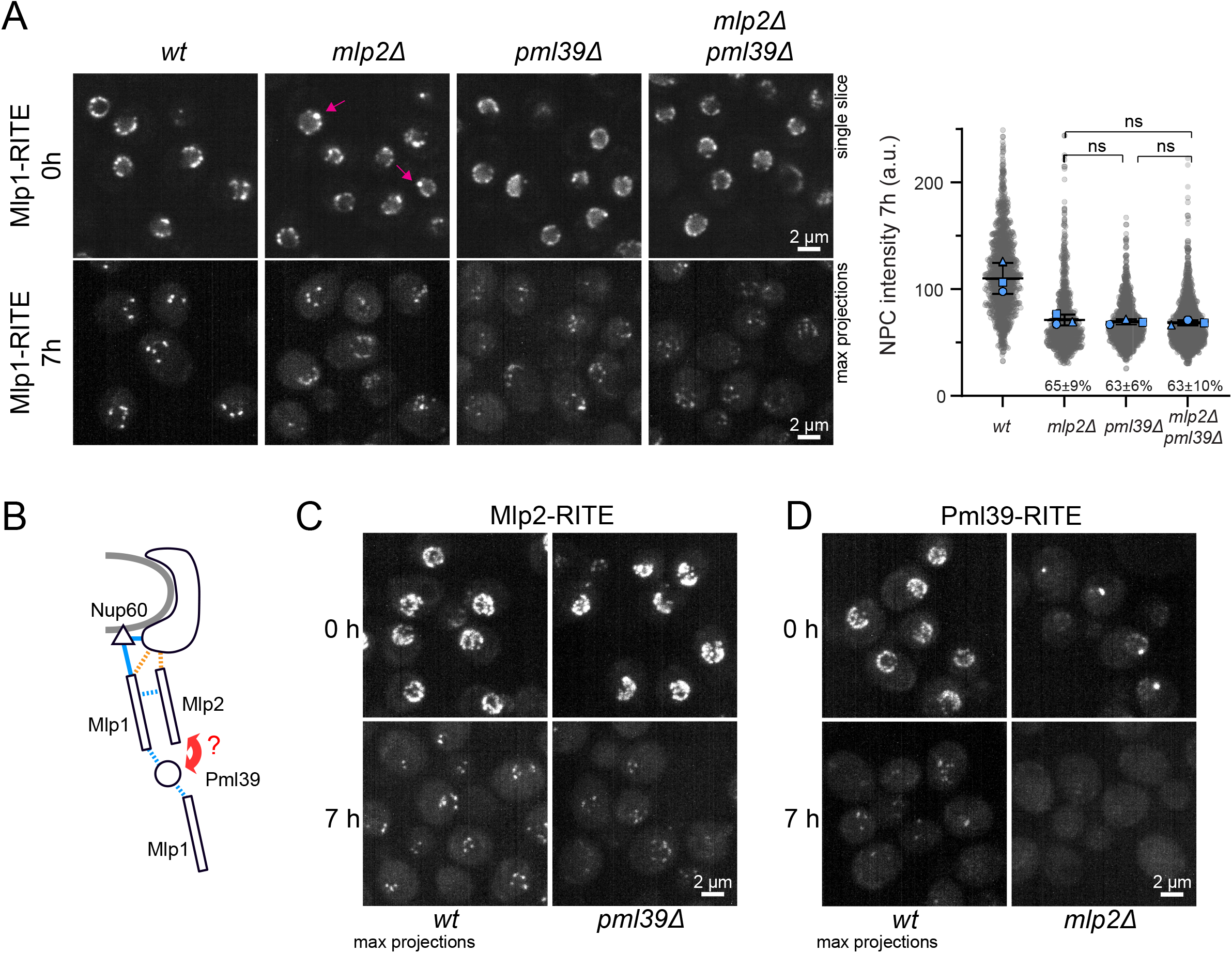
Mlp2 contributes to anchoring of Pml39 to the NPC. (A)doRITE assay of Mlp1 in different mutant backgrounds. Quantification on the right shows mean intensity measured for >70 NPCs at the 7 h timepoint per biological replicate and condition. Grey circles represent background-subtracted measurements from individual cells. Blue symbols represent means of individual biological replicates. Error bars represent the standard deviation. p values from paired two-tailed t test: ** < 0.01, * < 0.05, ns > 0.05.Quantification of intensity at the nuclear envelope at the 0h timepoint is shown in Fig. S4B.Pink arrows indicate Mlp1-EGFP foci which are prominent in the *mlp2Δ* strain. Quantification on the right shows mean intensity measured for >100 NPCs per biological replicate and condition at 7h after RITE induction. Grey circles represent background-subtracted measurements from individual cells. Blue symbols represent means of individual biological replicates. Error bars represent the standard deviation. Statistical analysis was carried out with a repeated measures one-way ANOVA with Tukey’s multiple comparison test. ns: not significant, * p< 0.05. All other comparison (wt with each mutant) are significant with p≤0.0001. Percentage values indicate the relative mean intensity compared to wt ± standard deviation. (B)Schematic illustration of the interaction analyzed in this figure: relationship between Mlp2 and Pml39. Dashed orange lines indicate interactions newly defined in the previous figures. (C)Cells expressing Mlp2-RITE without RITE induction or 7 hours after induction with or without Pml39 present. (D)Cells expressing Pml39-RITE without RITE induction or 7 hours after induction with or without Mlp2 present.

Because the reduction of Mlp1 at the NPC was reminiscent of the phenotype previously observed in *pml39Δ* cells (Gunkel et al., 2023; Zsok et al., 2024), we investigated whether these two proteins contribute to Mlp1 recruitment independently or via a shared pathway. We also carried out the doRITE experiments in *pml39Δ*, and *mlp2Δ pml39Δ* double deletion strains. All mutant backgrounds exhibited a similar reduction in Mlp1 intensity at the nuclear envelope in uninduced cells and at single NPC foci following doRITE induction (Figs 4A, S4A,B). However, there were distinct differences in the localization of unincorporated Mlp1. Prior to RITE induction, a subset of *mlp2Δ* cells exhibited nuclear Mlp1-EGFP foci, a phenotype typically seen when Mlp1 is prevented from efficiently assembling into NPCs and soluble copies are interlinked by Pml39 (Fig. 4A, pink arrows). Accordingly, both *pml39Δ* and *mlp2Δ pml39Δ* double mutant cells lacked these nuclear foci and instead displayed an increased pool of soluble nuclear Mlp1-EGFP (Fig. 4A). Together, these results suggest that Mlp2 and Pml39 operate in a common pathway to anchor the same extra pool of Mlp1 to the NPC.

Finally, we analyzed the reciprocal dependency of Pml39 and Mlp2 using doRITE (Fig. 4B). As shown previously (Gunkel et al., 2023), Mlp2 remained stably bound to NPCs in *pml39Δ* cells, though we noted a moderate reduction in intensity at individual NPCs (Figs 4C, S4C). In contrast, deletion of *mlp2* resulted in a severely compromised localization of Pml39, with no detectable signal at the 7-hour timepoint (Fig. 4D), indicating weak and unstable binding. Thus, Mlp2 is required for stable Pml39 recruitment.

### A three-way interaction network between Mlp1, Mlp2 and Pml39 is essential for constructing the nuclear basket

While Pml39 is known to interact with both Mlp1 and Mlp2 (Gunkel et al., 2023; Palancade et al., 2005), the molecular basis for these interactions remains unclear (Fig. 5A). To address this, we used AlphaFold3 (Abramson et al., 2024) to identify potential interaction interfaces within the Mlp1/Mlp2/Pml39 complex. Since Mlp1 and Mlp2 are thought to be constitutive dimers (Hase et al., 2001; Stankunas and Köhler, 2024), we modelled each as a homodimer. AlphaFold3 predicted interactions between Pml39 and each of the three previously identified interaction regions (Gunkel et al., 2023; Palancade et al., 2005): the Mlp1 N-terminus (aa1-120), the Mlp2 N-terminus (aa1-120) and the Mlp1 NBD (aa384-600) (Fig. 5B). The N-termini of Mlp1 and Mlp2 were predicted to bind the same cleft between the two lobes of Pml39, suggesting a mutually exclusive binding mode. The NBDs of Mlp1 was predicted to bind on the opposite side of Pml39, with terminal helices of Pml39 wrapping around the coiled-coil bundle. A similar interaction was also predicted for the NBD of Mlp2 (Fig. S5A), although there is no experimental evidence that supports such an interaction.

**Figure 5:**
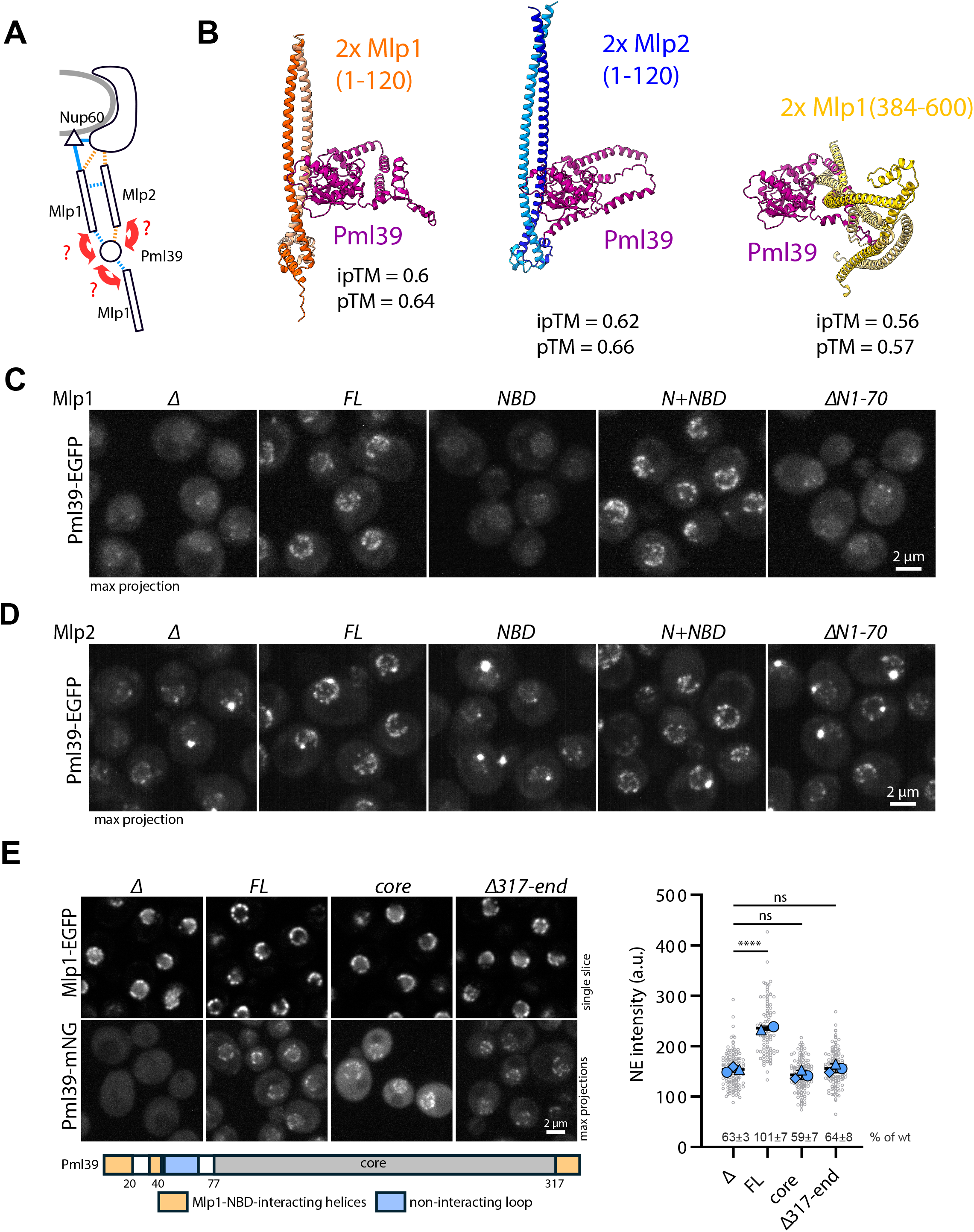
Investigating the interaction network of Pml39 at the NPC. (A)Schematic illustration of the interaction analyzed in this figure: Interaction of Pml39 with Mlp2 and two different regions of Mlp1. (B)AlphaFold3 predictions of Pml39 with dimers of Mlp1(aa 1-120), Mlp2(aa 1-120) or Mlp1 (aa 384-600). Pml39 is shown in the same orientation in all three complexes. (C)Localization of Pml39 upon rescue of mlp1Δ with different truncations of Mlp1. (D)Localization of Pml39 upon rescue of mlp2Δ with different truncations of Mlp2. (E)Localization of Mlp1-EGFP in Pml39 mutant strains. Bottom panels show the localization of the different Pml39 truncation mutants when tagged with mNG. Schematic indicates the different deleted regions on the Pml39 architecture. Plot shows quantification of Mlp1-EGFP intensity at the nuclear envelope. Grey circles represent background-subtracted measurements from individual cells (∼50 cells per condition and biological replicate). Blue symbols represent means of individual biological replicates. Error bars represent the standard deviation. Statistical testing was performed by fitting a mixed effects model with Sidak’s multiple comparison test. **** < 0.0001, ns > 0.05. Percentage values indicate the relative mean intensity compared to wt ± standard deviation.

To determine which of these interactions occur in the cell, we tested different truncations mutants. We first investigated the interaction between the N-termini of Mlp1 and Mlp2, which are likely responsible for recruiting Pml39 to the NPC. Upon deletion of Mlp1, Pml39 was lost from the NPC as previously reported (Gunkel et al., 2023; Palancade et al., 2005). Localization was rescued by re-introducing full-length Mlp1 or the N+NBD construct, but not by a construct lacking the N-terminus (Δ1-70) (Fig. 5C). This suggests that the Mlp1 N-terminus is necessary and sufficient for recruitment of Pml39 in the presence of Mlp2.

Very similarly, the N-terminus of Mlp2 was necessary and sufficient to promote recruitment of Pml39 to the NPC in the presence of Mlp1 (Fig. 5D), indicating that both N-termini cooperate to recruit Pml39. It is at present unclear how this occurs. When we modelled dimers of both N-termini and the Mlp1 NBD simultaneously in complex with Pml39, AlphaFold3 generated a high-confidence model in which the Mlp2 N-terminus occupied the primary binding site, while the Mlp1 N-terminus bound adjacent and parallel to it (Fig. S5B). The Mlp1 NBD occupied the same position as when modelled individually: binding to Pml39 on the opposite side from the Mlp1 and Mlp2 N-termini and perpendicular to them. Alternatively, it could also be that Mlp1 and Mlp2 form a heterodimer that is required to recruit Pml39. AlphaFold3 predicts complexes of Pml39 with heterodimers of the N-terminus and the NBDs with similar confidence values as for the homodimers (Fig. S5A).

### Pml39 terminal helices are required to anchor distal Mlp1 dimers

Finally, we investigated the mode by which Pml39 recruits additional copies of Mlp1 to the NPC. Our AlphaFold3 model predicts a distinct interaction interface between Pml39 and the Mlp1-NBD involving short alpha-helices at the N- and C-termini of Pml39 that wrap around the Mlp1-NBD bundle (Fig. 5B). We hypothesized that deleting the terminal helical regions of Pml39 would release the “distal” dimer of Mlp1 without affecting Pml39’s own localization. We therefore generated a truncated “core” Pml39 construct (Pml39core, aa 77-317) lacking the short N- and C-terminal extensions. Notably, this Pml39core construct retains some functionality of the full-length protein, since it was shown to rescue a synthetic growth phenotype of *nup133Δ* and *pml39Δ* (Hashimoto et al., 2022). In addition, we generated a set of truncation mutants lacking individual helices. We then tested whether these truncations were able to restore the full intensity of Mlp1 at the nuclear envelope when expressed in *pml39Δ* cells. All truncation mutants localized to the NPC, but only full-length Pml39 and a truncation lacking a loop region not predicted to interact with Mlp1 (Δ42-66) could rescue Mlp1 localization (Figs 5E, S5C). A partial rescue was observed with the construct lacking only the first 20 amin acids, all other truncation mutants failed to rescue Mlp1 localization. Although some of the truncation mutants exhibited increased expression and in particular increased cytoplasmic pools, this phenotype did not correlate with the ability to rescue Mlp1 localization. The overexpression phenotype appears to be caused by deletion of amino acids 20-40 and it could be that this region is involved either in targeting Pml39 to the nucleus or in promoting its degradation. Overall, these results validate the AlphaFold3 predictions and show that binding of Pml39 to the Mlp1 NBD requires the N- and C-termini of Pml39.

## Discussion

The results presented here allow us to propose a refined model for nuclear pore basket architecture. This model is consistent with a large number of observations from budding yeast and metazoa, and given the high level of conservation of the basket, it is likely that the organizational principles proposed here also apply to higher eukaryotes.

### Building a stable nuclear pore basket

Although Nup60 plays a central role in nuclear pore basket assembly by its ability to directly bind to the nuclear membrane, the nuclear outer ring, Mlp1 and Nup2 (Cibulka et al., 2022; Mészáros et al., 2015; Stankunas and Köhler, 2024), it is dispensable for maintenance of Mlp1 at the NPC at low temperatures or once basket assembly is completed (Aksenova et al., 2020; King et al., 2023; Ptak et al., 2025; Zsok et al., 2024). This implies the existence of so far unknown Nup60-independent interactions between Mlp1 and the nuclear pore scaffold. Our initial hypothesis - that the Mlp1 NBD is responsible for this scaffold contact - was based on current models for the interaction between Nup60, Mlp1 and the outer ring that were derived from structure predictions, interaction analyses and cryo-electron tomography (Akey et al., 2022; Singh et al., 2024; Stankunas and Köhler, 2024). However, this hypothesis proved not to be correct, and our data demonstrate that the NBD is insufficient for basket association in the absence of Nup60.

Instead, we found that Mlp1 and Mlp2 rely on each other to maintain association with the NPC in the absence of Nup60 (Fig. 3C-D). Lateral interactions between the coiled-coil regions likely combine with the interaction of each NBD with the NPC scaffold to cumulatively provide sufficient binding strength to maintain NPC association upon loss of Nup60.

### A stoichiometric model for nuclear basket architecture

By integrating our findings with measurements of Nup ratios (Zsok et al., 2024), we propose a stoichiometric model defined by a 4:2:1 ratio of Mlp1, Mlp2, and Pml39 (Fig. 6). In this model, the nuclear basket consists of 32 Mlp1 molecules, 16 Mlp2 molecules, and 8 Pml39 molecules. These numbers are best in line with the data we present and previous data. Mlp1 and Mlp2 are independently recruited to the NPC scaffold via interactions of their NBDs with Nup60 (Fig. 2B). Their independent recruitment supports the idea that each is recruited as a dimer. Extending from the NPC scaffold, they likely form a parallel helical bundle (Kim et al., 2018; Obarska-Kosinska et al., 2026 preprint). Independent scaffold binding of these two paralogues implicates the presence of two distinct binding sites. These might be provided by the presence of a “double” nuclear outer Y-ring, which was recently shown to occur on a subset of NPCs in budding yeast which also carry nuclear basket filaments (Akey et al., 2022; Singh et al., 2024).

**Figure 6:**
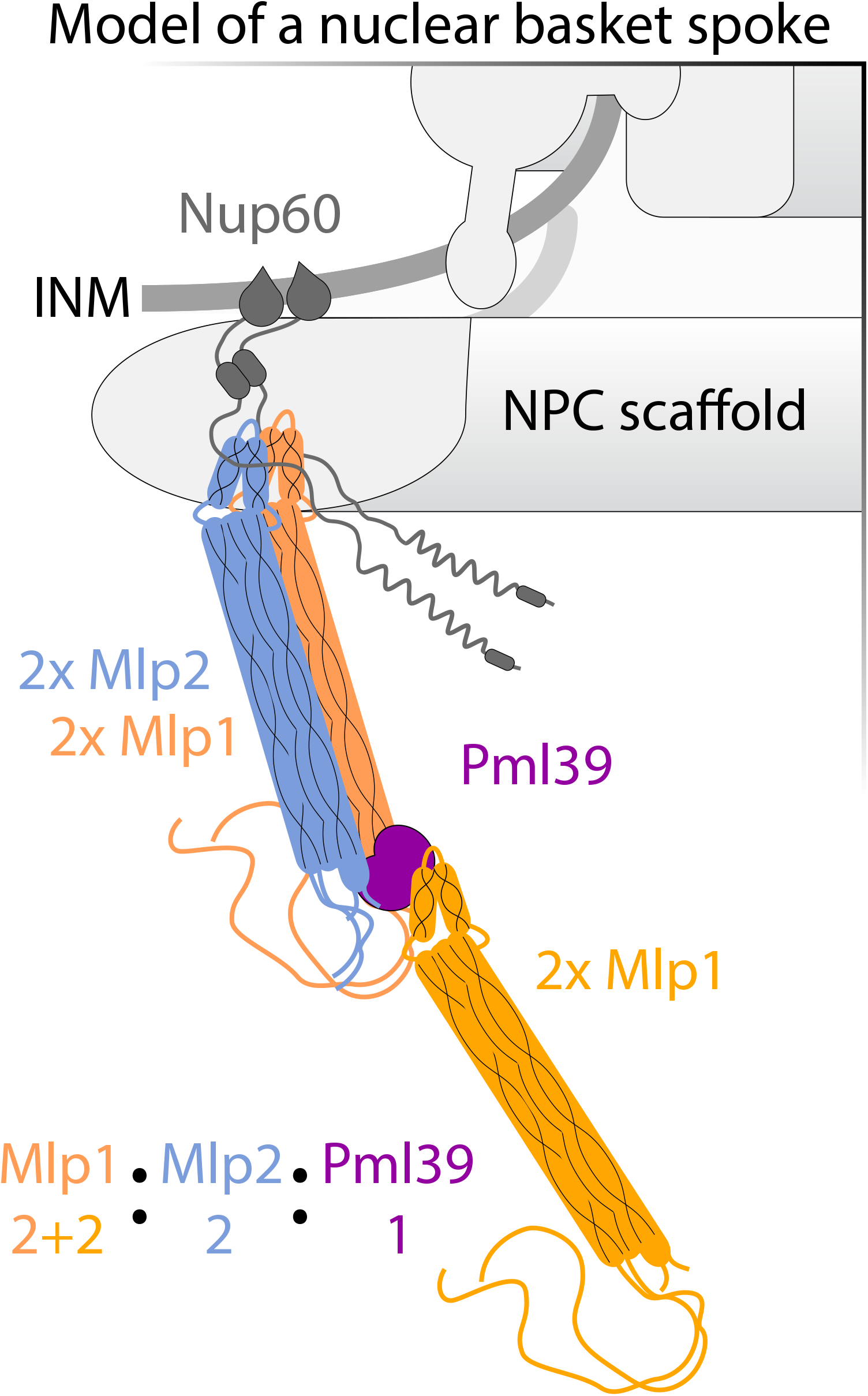
A model of nuclear pore basket architecture. Model for the architecture of one spoke of the nuclear pore basket. A dimer of Mlp1 and a dimer of Mlp2 are anchored to the NPC by Nup60 and via direct interactions with the scaffold. They form a parallel bundle. Pml39 is recruited via interaction with the N-termini of Mlp1 and Mlp2 dimers and in turn recruits another dimer of Mlp1 via its NPC binding domain. The stoichiometry of Mlp1:Mlp2:Pml39 at the NPC spoke is 4:2:1. Schematic based on (Dultz and Doye, 2025).

The N-termini of Mlp1 and Mlp2 extend into the nucleoplasm to create a composite binding site for Pml39. For efficient recruitment, both Mlp1 and Mlp2 are required, although weak and unstable recruitment is observed in the presence of Mlp1 alone (Fig. 4C, 5D). AlphaFold3 proposes a parallel orientation of the two homodimers, but an alternative model where Pml39 preferentially binds to a heterodimer of Mlp1 and Mlp2 also satisfies the requirement that both be present. Once anchored, Pml39 utilizes an independent surface on its opposite face to recruit an additional Mlp1 dimer via its NBD. Although the reduction of Mlp1 at the nuclear envelope upon loss of Pml39 is apparently less than 50 % in our and previous quantifications (Fig. 4A, (Gunkel et al., 2023; Zsok et al., 2024), this might be due inaccureate background subtraction caused by the high level of nucleoplasmic background in these cells.

### Evolutionary conservation of nuclear basket architecture

Our model aligns closely with recent models of the metazoan NPC (Obarska-Kosinska et al., 2026 preprint), which based on EM structures propose that two TPR dimers bind to the nuclear outer ring with a third TPR dimer recruited via ZC3HC1. However, a discrepancy remains regarding stoichiometry. In both systems, the loss of Pml39/ZC3HC1 reduces Mlp1/TPR intensity by approximately 50% (Gunkel et al., 2023, 2021; Zsok et al., 2024). While this fits our yeast model, it suggests that in metazoa, ZC3HC1 might recruit more than one TPR dimer, or that the baseline number of TPR dimers is different.

The conformation of the distal TPR/Mlp1 dimer remains an open question. While the recent cryo-electron tomography maps indicate that the first layer of TPR/Mlp1/2 may form rigid rod-like structures (Obarska-Kosinska et al., 2026 preprint; Singh et al., 2024), the conformation of the distal layer remains unclear. It could interlink the filaments of adjacent spokes to form the distal ring (Goldberg and Allen, 1992; Kiseleva et al., 1998) or extend further into the nucleoplasm to extend the basket filaments. The latter is supported by the observation of “basket ladders” in Xenopus oocytes, which were proposed to form by consecutive alternate recruitment of ZC3HC1 and TPR (Gunkel et al., 2021). However, we find that the recruitment of Pml39 requires both Mlp1 and Mlp2 N-termini, while only Mlp1 is recruited in the “distal” layer. This would preclude the formation of basket ladders and suggest that a strictly stoichiometric filament organization is favored in yeast.

The metazoan models also indicate - in agreement with previous functional data (Aksenova et al., 2020; Umlauf et al., 2013) - that the TREX-2 complex is an integral structural component of the nuclear pore basket and binds both the outer ring and the NBD of TPR. It is plausible that this could also be the case in budding yeast.

### Conclusion

In conclusion, we propose a refined nuclear pore basket model that is fully consistent with previous results characterizing the interactions between basket Nups in both yeast and metazoa. Nevertheless, several questions regarding the architecture and the organization of the nuclear basket remain. This includes the exact stoichiometry of the basket components, the conformation of the proximal and in particular the distal Mlp1/2 dimers, the composition of the distal ring and the detailed interaction between the basket filaments and the scaffold. Our refined model will help to inform future work to address these questions.

## Limitations

Although all our data are consistent with the presented model, alternative explanations cannot be excluded. In particular, it remains unclear how the N-termini of Mlp1 and Mlp2 interact to stably bind Pml39. Furthermore, our quantitative fluorescence microscopy measurements have some limitations. First, due to the high NPC density in yeast we can only analyze the overall intensity at the nuclear envelope and cannot measure intensities at individual NPCs. Therefore, we cannot differentiate between direct NPC association and potential, additional, nuclear envelope associated pools. Furthermore, limitations apply to the comparison of intensity measurements carried out in cells that have different levels in soluble background. In particular, this applies to strains that express different Mlp1 truncations or our measurements of Mlp1 in *pml39Δ* strains that exhibit an increased nucleoplasmic pool. Our measurements in cells where only NPCs older than 7 hours are measured are not affected by this, but in this case differences in a potential slow exchange of individual NPC subunits could affect the observed comparisons.

## Materials and Methods

### Plasmids and yeast strains

Plasmids were generated using restriction enzyme based cloning or Golden Gate Assembly (Lee et al., 2015). Inserts of genomic sequences into Golden Gate assembly vectors were generated either by PCR on genomic DNA or, if codons had to be optimized to remove internal restriction sites, ordered from Twist Bioscience (South San Francisco, USA). Yeast strains were constructed by integration of linearized plasmids or PCR products with homologous ends targeting integration (Longtine et al., 1998). Some strains with multiple genetic modifications were generated by mating, sporulation and spore selection. Tables of the used strains and plasmids can be found in Tables S1 and S2. Strains and plasmids are available to the research community upon reasonable request.

The nuclear localization sequences of Mlp1 and Mlp2 are located in their C-terminal region (Strambio-de-Castillia et al., 1999). Since many of our truncation constructs thus lack an NLS, we added an SV40NLS in most constructs of Mlp1 and Mlp2 including the respective full-length controls. The presence of an exogenous NLS is in each case indicated in the strain information in Table S1.

### Yeast culture conditions and treatments

Yeast cultures were grown in synthetic medium containing 2% glucose at 30°C unless otherwise stated. All experiments were carried out with cells growing exponentially. Prior to induction of recombination, RITE strains were maintained and grown on selective (-ura) solid or liquid medium to select against a low level of uninduced recombination from the Cre recombinase which is continuously expressed under control of a TDH3 promoter (Terweij et al., 2013).

Exponentially growing cells were induced to recombine with 0.5 µM β-estradiol (Sigma), and cultures were imaged in exponential growth phase after the indicated times. Auxin-inducible degradation of proteins was induced by treatment of exponentially growing cells with 0.5 mM indole-3-acetic acid (IAA) (Sigma, I2886-5G, CAS: 87-51-4) and 4 µM phytic acid dipotassium salt (Sigma, 5681, CAS: 129832-03-7) for the indicated time.

### Live cell microscopy

Cells were imaged in 384-well plates (Matrical) coated with concanavalin A for immobilization. Three-dimensional volume stacks of yeast cells were acquired on a temperature-controlled inverted Nipkow spinning disk microscope equipped with the Yokogawa Confocal Scanner Unit CSU-W1-T2 SoRa and a triggered Piezo z-stage (Mad City Labs Nano-Drive). It was used in spinning disk mode with a pinhole diameter of 50 µm combined with a 1.45 NA, 100x objective and controlled by the NIS Elements Software (Nikon). Images were acquired with a sCMOS Hamamatsu Orca Fusion BT camera (2304 × 2304 pixel, 6.5 × 6.5 µm pixel size). Imaging was performed at the same temperature as growth incubation with 50% laser intensity of a Oxxius 488 nm 200 mW LNC Version light source. Z-stack data were taken with the Piezo stage in triggering mode with 0.2 µm sectioning. Exposure times were 200-500 ms. In general, all experiments were carried out in at least three biological replicates on different days.

### Western blots

Logarithmically growing cells were lysed with 0.1M NaOH for 15 min at RT and subsequently boiled in NuPAGE 1xSDS loading buffer supplemented with 50 mM DTT at 95°C for 5 min. An amount corresponding to 1 OD of cells was loaded per lane and separated on NuPAGE 4–12% Bis–Tris 1.0 mm miniprotein gels (Thermo Fisher, NP0321BOX) with 200 V for 60 min. Proteins were transferred to 0.2 µM nitrocellulose membranes (Amersham™ Protran®, GE10600002) with a Trans-Blot® Turbo™ semi-dry transfer system (Bio-Rad) at 25 V. The transfer was verified by a Ponceau-S quick protein stain. Membranes were blocked in PBS containing 5% (w/v) milk for at least 1.5 h at room temperature and incubated with primary antibodies diluted in the same buffer overnight at 4°C. Membranes were then washed three times for 10 minutes each with 1xPBS and incubated with secondary antibodies that were diluted in 1xPBS containing 5%milk for at least 1.5 h at room temperature in the dark. This was followed by three additional 10 min washes with 1xPBS. The membranes were imaged using an Odyssey CLx imaging system (LI-COR Biosciences, Lincoln, NE). The following antibodies were used: Mouse anti-V5 tag (1:2000; Bio-Rad Cat# MCA1360, RRID:AB_322378), Rabbit anti-Crm1 (1:5000) (Zeitler and Weis, 2004), IRDye® 800CW Goat anti-Rabbit IgG (H + L) (1:10’000; LICORbio Cat# 926-32211, RRID:AB_621843), Alexa 680 Goat anti-Mouse (1:10’000; Thermo Fisher Scientific Cat# A-21057, RRID:AB_2535723). Figure preparation and western blot quantifications were done using Fiji/ImageJ and Image Studio™ software version 6.0(LI-COR Biosciences). Uncropped blots are shown in Fig. S6.

### Image and data analysis

Image analysis and adjustments were carried out in FIJI (Schindelin et al., 2012). Data analysis, curve fitting, statistical analysis, and plotting were carried out in Excel or GraphPad Prism. Analyzed cell numbers are listed in Table S3 if not specified elsewhere.

For nuclear envelope intensity measurements, at least 50 nuclei per condition were analyzed in three biological replicates. A hand-drawn line of width 3 pixels following the NE as delineated by the Nup-EGFP signal was created in FIJI omitting the nucleolar region identified by a gap in Nup localization. The mean of nucleoplasmic and cytoplasmic background was subtracted.

Intensities of NPCs 7h after RITE induction were determined on maxima identified by the “Find Maxima…” tool in FIJI on maximum projections using a prominence of 20 as the cut-off. For each detected maximum, the brightest slice was then selected and the mean intensity in a refined circular region of interest of four pixel diameter was determined. Background was subtracted. Measurements were filtered for datapoints with low contrast.

Total cell intensities were measured by segmenting cells on an image filtered with a rolling ball filter in FIJI via thresholding and subtracting extracellular background from the integrated intensity. Cells not expressing fluorescent proteins were used to determine the background from autofluorescence.

### AlphaFold predictions

Structure predictions were carried out with AlphaFold3 on the Google Server (Abramson et al., 2024). Structural representations in Figs 5 and S5 were generated with ChimeraX.

### Generative AI

Gemini 3 was used to improve text, spelling and grammar. All text was checked by the authors, and the authors take full responsibility for the content.

## Author Contributions

KS: Conceptualization, Methodology, Investigation, Formal analysis, Data curation, Writing – review and editing, Validation, Visualization. AS: Conceptualization, Methodology, Investigation, Formal analysis, Data curation, Writing – review and editing, Validation, Visualization. ED: Conceptualization, Methodology, Investigation, Formal analysis, Data curation, Writing - original draft, Validation, Visualization, Supervision, Project administration, Funding acquisition.

### Conflicts of interest

The authors declare no financial conflict of interest.

## Acknowledgements

We are grateful to Karsten Weis, Jonas Fischer and Sarah Khawaja as well as members of the Weis lab for discussions and critical reading of the manuscript. We thank ScopeM for excellent microscopy support. This work was supported by an SNSF Research Grant (320030-236124 to E.D.).

## Abbreviations

NPC: nuclear pore complex
mNG: monomeric neon green
EGFP: enhanced green fluorescent protein
NLS: nuclear localization signal
Nup: nucleoporin

## Notes

### Competing Interest Statement

The authors have declared no competing interest.

### Summary of Updates

additional data added: - Mlp1 and Mlp2 depend on each other to remain associated with the NPC upon depletion of Nup60 - Mlp2 relies on Mlp1 to stabilize its interaction with the NPC through multiple generations - Pml39 relies on the N-termini of both Mlp1 and Mlp2 for recruitment to the NPC - all the N- and C-terminal helices of Pml39 are required to recruit additional Mlp1 to the NPC

## References

Abramson, J., Adler, J., Dunger, J., Evans, R., Green, T., Pritzel, A., Ronneberger, O., Willmore, L., Ballard, A.J., Bambrick, J., Bodenstein, S.W., Evans, D.A., Hung, C.-C., O’Neill, M., Reiman, D., Tunyasuvunakool, K., Wu, Z., Žemgulytė, A., Arvaniti, E., Beattie, C., Bertolli, O., Bridgland, A., Cherepanov, A., Congreve, M., Cowen-Rivers, A.I., Cowie, A., Figurnov, M., Fuchs, F.B., Gladman, H., Jain, R., Khan, Y.A., Low, C.M.R., Perlin, K., Potapenko, A., Savy, P., Singh, S., Stecula, A., Thillaisundaram, A., Tong, C., Yakneen, S., Zhong, E.D., Zielinski, M., Žídek, A., Bapst, V., Kohli, P., Jaderberg, M., Hassabis, D., Jumper, J.M., 2024. Accurate structure prediction of biomolecular interactions with AlphaFold 3. Nature 630, 493–500. 10.1038/s41586-024-07487-w

Akey, C.W., Singh, D., Ouch, C., Echeverria, I., Nudelman, I., Varberg, J.M., Yu, Z., Fang, F., Shi, Y., Wang, J., Salzberg, D., Song, K., Xu, C., Gumbart, J.C., Suslov, S., Unruh, J., Jaspersen, S.L., Chait, B.T., Sali, A., Fernandez-Martinez, J., Ludtke, S.J., Villa, E., Rout, M.P., 2022. Comprehensive structure and functional adaptations of the yeast nuclear pore complex. Cell 185, 361–378 e25. 10.1016/j.cell.2021.12.015

Aksenova, V., Smith, A., Lee, H., Bhat, P., Esnault, C., Chen, S., Iben, J., Kaufhold, R., Yau, K.C., Echeverria, C., Fontoura, B., Arnaoutov, A., Dasso, M., 2020. Nucleoporin TPR is an integral component of the TREX-2 mRNA export pathway. Nat Commun 11, 4577. 10.1038/s41467-020-18266-2

Bensidoun, P., Reiter, T., Montpetit, B., Zenklusen, D., Oeffinger, M., 2022. Nuclear mRNA metabolism drives selective basket assembly on a subset of nuclear pore complexes in budding yeast. Mol. Cell 82, 3856–3871.e6. 10.1016/j.molcel.2022.09.019

Camacho, C., Coulouris, G., Avagyan, V., Ma, N., Papadopoulos, J., Bealer, K., Madden, T.L., 2009. BLAST+: architecture and applications. BMC Bioinformatics 10, 421. 10.1186/1471-2105-10-421

Cibulka, J., Bisaccia, F., Radisavljevic, K., Gudino Carrillo, R.M., Kohler, A., 2022. Assembly principle of a membrane-anchored nuclear pore basket scaffold. Sci Adv 8, eabl6863. 10.1126/sciadv.abl6863

De Magistris, P., 2021. The Great Escape: mRNA Export through the Nuclear Pore Complex. Int. J. Mol. Sci. 22, 11767. 10.3390/ijms222111767

Dultz, E., Doye, V., 2025. Opening the gate: Complexity and modularity of the nuclear pore scaffold and basket. Curr. Opin. Cell Biol. 92, 102461. 10.1016/j.ceb.2024.102461

Fasken, M.B., Stewart, M., Corbett, A.H., 2008. Functional significance of the interaction between the mRNA-binding protein, Nab2, and the nuclear pore-associated protein, Mlp1, in mRNA export. J. Biol. Chem. 283, 27130–27143. 10.1074/jbc.M803649200

Feuerbach, F., Galy, V., Trelles-Sticken, E., Fromont-Racine, M., Jacquier, A., Gilson, E., Olivo-Marin, J.-C., Scherthan, H., Nehrbass, U., 2002. Nuclear architecture and spatial positioning help establish transcriptional states of telomeres in yeast. Nat. Cell Biol. 4, 214–221. 10.1038/ncb756

Gabler, F., Nam, S.-Z., Till, S., Mirdita, M., Steinegger, M., Söding, J., Lupas, A.N., Alva, V., 2020. Protein Sequence Analysis Using the MPI Bioinformatics Toolkit. Curr. Protoc. Bioinforma. 72, e108. 10.1002/cpbi.108

Galy, V., Gadal, O., Fromont-Racine, M., Romano, A., Jacquier, A., Nehrbass, U., 2004. Nuclear retention of unspliced mRNAs in yeast is mediated by perinuclear Mlp1. Cell 116, 63–73.

Goldberg, M.W., Allen, T.D., 1992. High resolution scanning electron microscopy of the nuclear envelope: demonstration of a new, regular, fibrous lattice attached to the baskets of the nucleoplasmic face of the nuclear pores. J. Cell Biol. 119, 1429–1440. 10.1083/jcb.119.6.1429

Gunkel, P., Cordes, V.C., 2022. ZC3HC1 is a structural element of the nuclear basket effecting interlinkage of TPR polypeptides. Mol. Biol. Cell 33, ar82. 10.1091/mbc.E22-02-0037

Gunkel, P., Iino, H., Krull, S., Cordes, V.C., 2023. An evolutionarily conserved bimodular domain anchors ZC3HC1 and its yeast homologue Pml39p to the nuclear basket. Mol. Biol. Cell 34, ar40. 10.1091/mbc.E22-09-0402

Gunkel, P., Iino, H., Krull, S., Cordes, V.C., 2021. ZC3HC1 Is a Novel Inherent Component of the Nuclear Basket, Resident in a State of Reciprocal Dependence with TPR. Cells 10. 10.3390/cells10081937

Hakhverdyan, Z., Molloy, K.R., Keegan, S., Herricks, T., Lepore, D.M., Munson, M., Subbotin, R.I., Fenyo, D., Aitchison, J.D., Fernandez-Martinez, J., Chait, B.T., Rout, M.P., 2021. Dissecting the Structural Dynamics of the Nuclear Pore Complex. Mol. Cell 81, 153–165 e7. 10.1016/j.molcel.2020.11.032

Hase, M.E., Kuznetsov, N.V., Cordes, V.C., 2001. Amino Acid Substitutions of Coiled-Coil Protein Tpr Abrogate Anchorage to the Nuclear Pore Complex but Not Parallel, In-Register Homodimerization. Mol. Biol. Cell 12, 2433. 10.1091/mbc.12.8.2433

Hashimoto, H., Ramirez, D.H., Lautier, O., Pawlak, N., Blobel, G., Palancade, B., Debler, E.W., 2022. Structure of the pre-mRNA leakage 39-kDa protein reveals a single domain of integrated zf-C3HC and Rsm1 modules. Sci. Rep. 12, 17691. 10.1038/s41598-022-22183-3

Huang, G., Zeng, C., Shi, Y., 2023. Structure of the nuclear pore complex goes atomic. Curr. Opin. Struct. Biol. 78, 102523. 10.1016/j.sbi.2022.102523

Kim, S.J., Fernandez-Martinez, J., Nudelman, I., Shi, Y., Zhang, W., Raveh, B., Herricks, T., Slaughter, B.D., Hogan, J.A., Upla, P., Chemmama, I.E., Pellarin, R., Echeverria, I., Shivaraju, M., Chaudhury, A.S., Wang, J., Williams, R., Unruh, J.R., Greenberg, C.H., Jacobs, E.Y., Yu, Z., de la Cruz, M.J., Mironska, R., Stokes, D.L., Aitchison, J.D., Jarrold, M.F., Gerton, J.L., Ludtke, S.J., Akey, C.W., Chait, B.T., Sali, A., Rout, M.P., 2018. Integrative structure and functional anatomy of a nuclear pore complex. Nature 555, 475–482. 10.1038/nature26003

King, G.A., Wettstein, R., Varberg, J.M., Chetlapalli, K., Walsh, M.E., Gillet, L.C.J., Hernández-Armenta, C., Beltrao, P., Aebersold, R., Jaspersen, S.L., Matos, J., Ünal, E., 2023. Meiotic nuclear pore complex remodeling provides key insights into nuclear basket organization. J. Cell Biol. 222, e202204039. 10.1083/jcb.202204039

Kiseleva, E., Goldberg, M.W., Allen, T.D., Akey, C.W., 1998. Active nuclear pore complexes in Chironomus: visualization of transporter configurations related to mRNP export. J. Cell Sci. 111 (Pt 2), 223–236. 10.1242/jcs.111.2.223

Krull, S., Thyberg, J., Bjorkroth, B., Rackwitz, H.R., Cordes, V.C., 2004. Nucleoporins as components of the nuclear pore complex core structure and Tpr as the architectural element of the nuclear basket. Mol Biol Cell 15, 4261–77. 10.1091/mbc.e04-03-0165

Lee, M.E., DeLoache, W.C., Cervantes, B., Dueber, J.E., 2015. A Highly Characterized Yeast Toolkit for Modular, Multipart Assembly. ACS Synth. Biol. 4, 975–986. 10.1021/sb500366v

Longtine, M.S., McKenzie, A., Demarini, D.J., Shah, N.G., Wach, A., Brachat, A., Philippsen, P., Pringle, J.R., 1998. Additional modules for versatile and economical PCR-based gene deletion and modification in Saccharomyces cerevisiae. Yeast Chichester Engl. 14, 953–961. 10.1002/(SICI)1097-0061(199807)14:10%3C953::AID-YEA293%3E3.0.CO;2-U

McNeil, J.B., Lee, S.-K., Oliinyk, A., Raina, S., Garg, J., Moallem, M., Urquhart-Cox, V., Fillingham, J., Cheung, P., Rosonina, E., 2024. 1,10-phenanthroline inhibits sumoylation and reveals that yeast SUMO modifications are highly transient. EMBO Rep. 25, 68–81. 10.1038/s44319-023-00010-8

Mészáros, N., Cibulka, J., Mendiburo, M.J., Romanauska, A., Schneider, M., Köhler, A., 2015. Nuclear pore basket proteins are tethered to the nuclear envelope and can regulate membrane curvature. Dev. Cell 33, 285–298. 10.1016/j.devcel.2015.02.017

Niepel, M., Molloy, K.R., Williams, R., Farr, J.C., Meinema, A.C., Vecchietti, N., Cristea, I.M., Chait, B.T., Rout, M.P., Strambio-De-Castillia, C., 2013. The nuclear basket proteins Mlp1p and Mlp2p are part of a dynamic interactome including Esc1p and the proteasome. Mol. Biol. Cell 24, 3920–3938. 10.1091/mbc.E13-07-0412

Nishimura, K., Fukagawa, T., Takisawa, H., Kakimoto, T., Kanemaki, M., 2009. An auxin-based degron system for the rapid depletion of proteins in nonplant cells. Nat Methods 6, 917–22. 10.1038/nmeth.1401

Nobari, P., Doye, V., Boumendil, C., 2023. Metazoan nuclear pore complexes in gene regulation and genome stability. DNA Repair 130, 103565. 10.1016/j.dnarep.2023.103565

Obarska-Kosinska, A., Zhu, Y., Geißler, K., Rosenkranz, R.R.E., Yokoyama, N., Kreysing, J.P., Xing, H., Glushkova, D., Kubańska, M.A., Böhm, S., Kräusslich, H.-G., Turoňová, B., Liu, F., Beck, M., 2026. How the TREX-2 complex associates with the nuclear pore. preprint bioRxiv 10.64898/2026.02.20.707084

Onischenko, E., Noor, E., Fischer, J.S., Gillet, L., Wojtynek, M., Vallotton, P., Weis, K., 2020. Maturation Kinetics of a Multiprotein Complex Revealed by Metabolic Labeling. Cell 183, 1785–1800 e26. 10.1016/j.cell.2020.11.001

Palancade, B., Zuccolo, M., Loeillet, S., Nicolas, A., Doye, V., 2005. Pml39, a novel protein of the nuclear periphery required for nuclear retention of improper messenger ribonucleoparticles. Mol Biol Cell 16, 5258–68. 10.1091/mbc.e05-06-0527

Petrovic, S., Mobbs, G.W., Bley, C.J., Nie, S., Patke, A., Hoelz, A., 2022. Structure and Function of the Nuclear Pore Complex. Cold Spring Harb. Perspect. Biol. 14, a041264. 10.1101/cshperspect.a041264

Ptak, C., Saik, N.O., Wozniak, R.W., 2025. Ulp1 association with nuclear pore complexes is required for the maintenance of global SUMOylation. Mol. Biol. Cell 36, ar81. 10.1091/mbc.E24-12-0563

Schindelin, J., Arganda-Carreras, I., Frise, E., Kaynig, V., Longair, M., Pietzsch, T., Preibisch, S., Rueden, C., Saalfeld, S., Schmid, B., Tinevez, J.-Y., White, D.J., Hartenstein, V., Eliceiri, K., Tomancak, P., Cardona, A., 2012. Fiji: an open-source platform for biological-image analysis. Nat. Methods 9, 676–682. 10.1038/nmeth.2019

Simon, M.-N., Dubrana, K., Palancade, B., 2024. On the edge: how nuclear pore complexes rule genome stability. Curr. Opin. Genet. Dev. 84, 102150. 10.1016/j.gde.2023.102150

Singh, D., Soni, N., Hutchings, J., Echeverria, I., Shaikh, F., Duquette, M., Suslov, S., Li, Z., van Eeuwen, T., Molloy, K., Shi, Y., Wang, J., Guo, Q., Chait, B.T., Fernandez-Martinez, J., Rout, M.P., Sali, A., Villa, E., 2024. The molecular architecture of the nuclear basket. Cell S0092-8674(24)00780–3. 10.1016/j.cell.2024.07.020

Stankunas, E., Köhler, A., 2024. Docking a flexible basket onto the core of the nuclear pore complex. Nat. Cell Biol. 26, 1504–1519. 10.1038/s41556-024-01484-x

Strambio-de-Castillia, C., Blobel, G., Rout, M.P., 1999. Proteins connecting the nuclear pore complex with the nuclear interior. J. Cell Biol. 144, 839–855. 10.1083/jcb.144.5.839

Terweij, M., van Welsem, T., van Deventer, S., Verzijlbergen, K.F., Menendez-Benito, V., Ontoso, D., San-Segundo, P., Neefjes, J., van Leeuwen, F., 2013. Recombination-induced tag exchange (RITE) cassette series to monitor protein dynamics in Saccharomyces cerevisiae. G3 Bethesda 3, 1261–72. 10.1534/g3.113.006213

Umlauf, D., Bonnet, J., Waharte, F., Fournier, M., Stierle, M., Fischer, B., Brino, L., Devys, D., Tora, L., 2013. The human TREX-2 complex is stably associated with the nuclear pore basket. J. Cell Sci. 126, 2656–2667. 10.1242/jcs.118000

Zeitler, B., Weis, K., 2004. The FG-repeat asymmetry of the nuclear pore complex is dispensable for bulk nucleocytoplasmic transport in vivo. J. Cell Biol. 167, 583–590. 10.1083/jcb.200407156

Zsok, J., Simon, F., Bayrak, G., Isaki, L., Kerff, N., Kicheva, Y., Wolstenholme, A., Weiss, L.E., Dultz, E., 2024. Nuclear basket proteins regulate the distribution and mobility of nuclear pore complexes in budding yeast. 10.1101/2023.09.28.558499

