## Supplemental Material for "Mlp1 and Mlp2 cooperate to build a stoichiometric nuclear pore basket in budding yeast"

### Figure S1

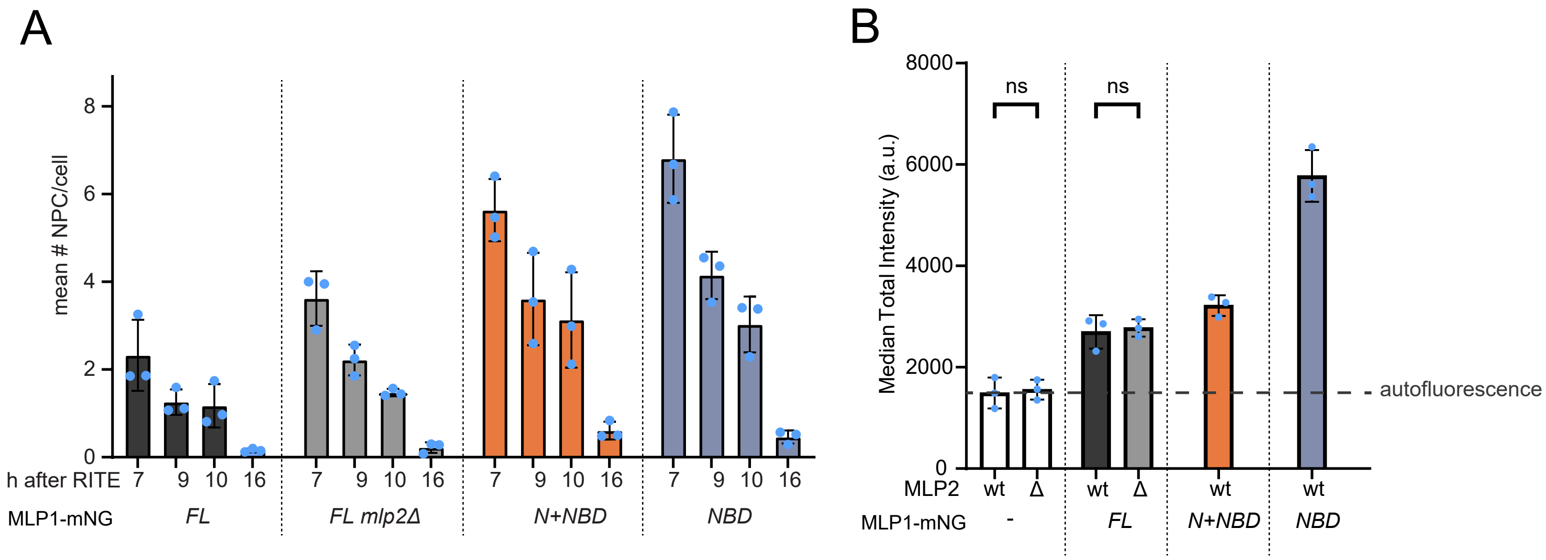


**Figure S1: Mlp1 truncations bind the NPC stably.**
(A) Quantification of foci (single NPCs) in doRITE assay with Mlp1 truncation constructs (compare Figure 1D). Dots were manually counted in z stacks of 100 randomly chosen cells per time point and replicate. Blue dots represent biological replicates. Error bars represent the standard deviation.
(B) Total fluorescence intensity of cells shown in Figure 1D at the 0 h timepoint. Blue dots represent biological replicates. Error bars represent the standard deviation. wt and *mlp2Δ* were not significantly different by paired two-tailed t-test (ns).

### Figure S2

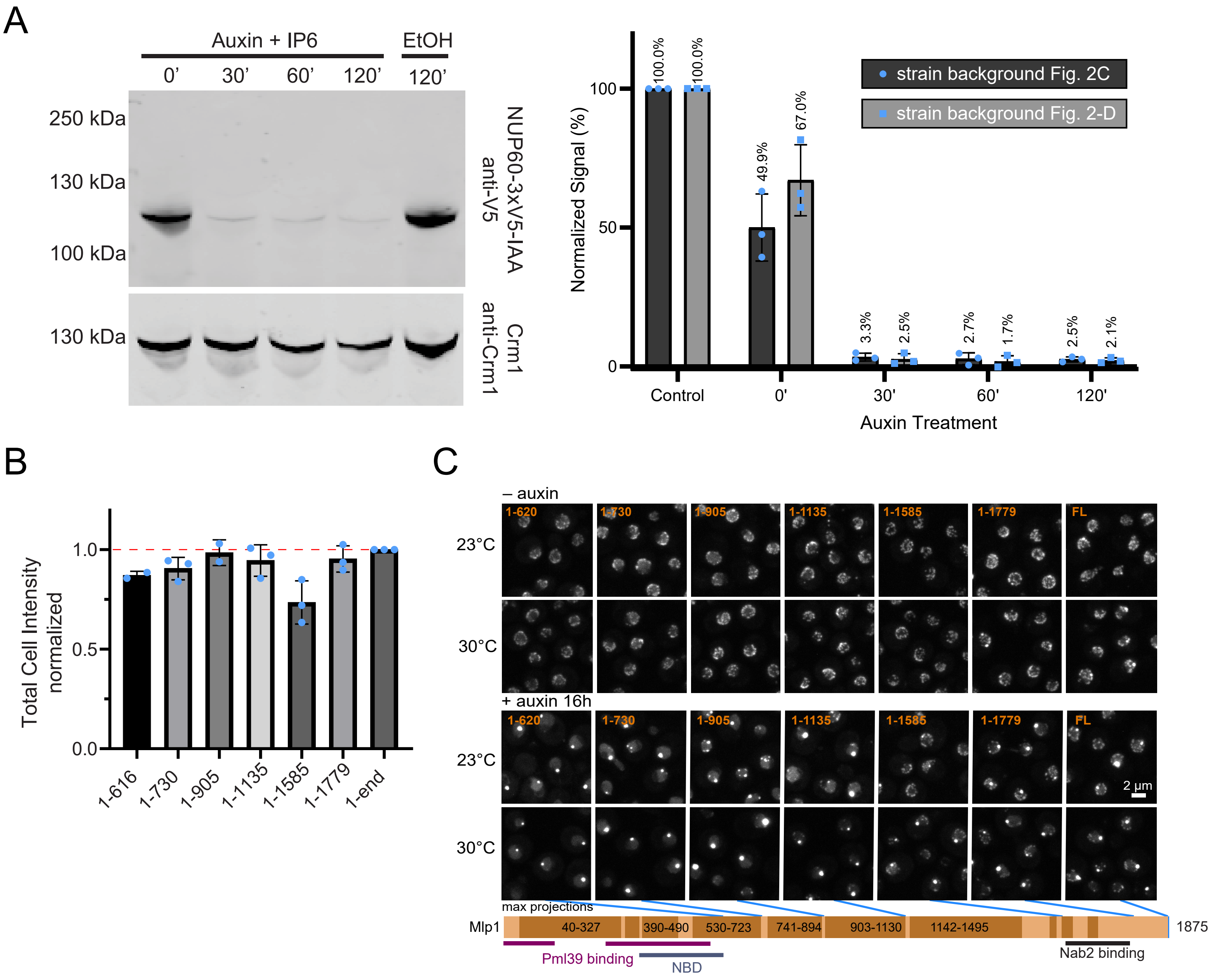


**Figure S2: The Mlp1 C-terminus confers Nup60-independent binding at the NPC.**
(A) Western blot showing depletion of Nup60 upon addition of auxin. Example blot of the strain background for the strains in Figure 2C is shown. Quantification on the right shows biological triplicates for the two strain backgrounds used in Figure 2C and 2D&S2B.
(B) Total fluorescence intensity of cells expressing different C-terminal truncations of Mlp1 in the absence of auxin to compare expression levels. Total fluorescence intensity was normalized to the full-length construct. Blue dots represent biological replicates. Bars show the mean, error bars represent the standard deviation.
(C) Localization of NLS-EGFP tagged, C-terminally truncated versions of Mlp1 after overnight growth in the absence of Nup60 by auxin-induced depletion at either 23 °C or 30 °C.
Image panels shown are maximum intensity projections.

### Figure S3

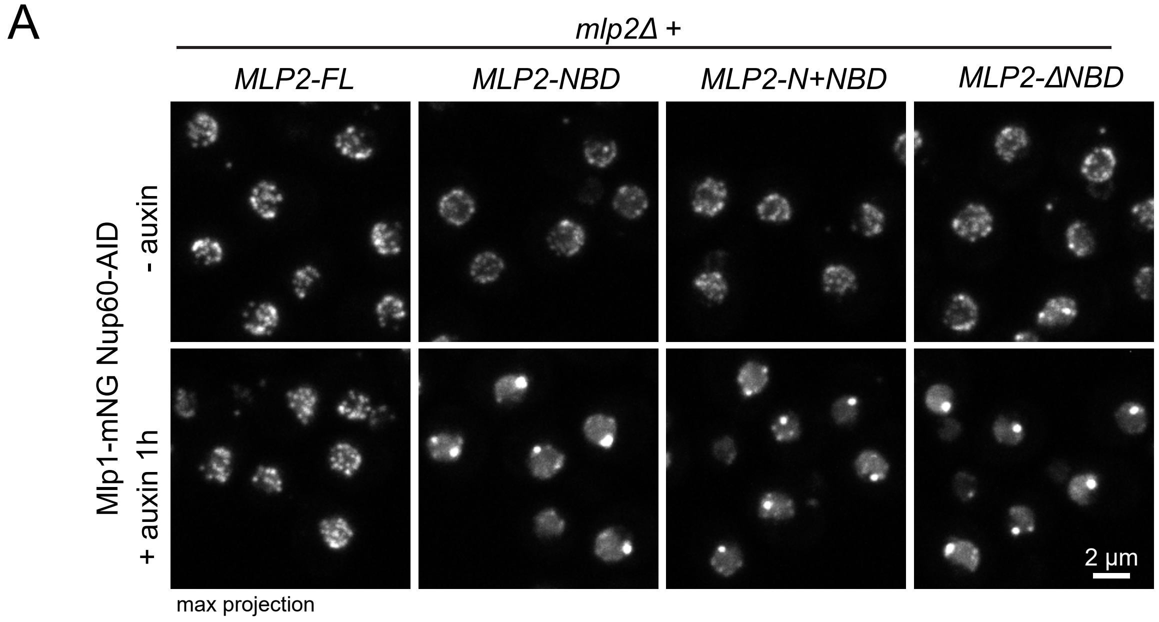


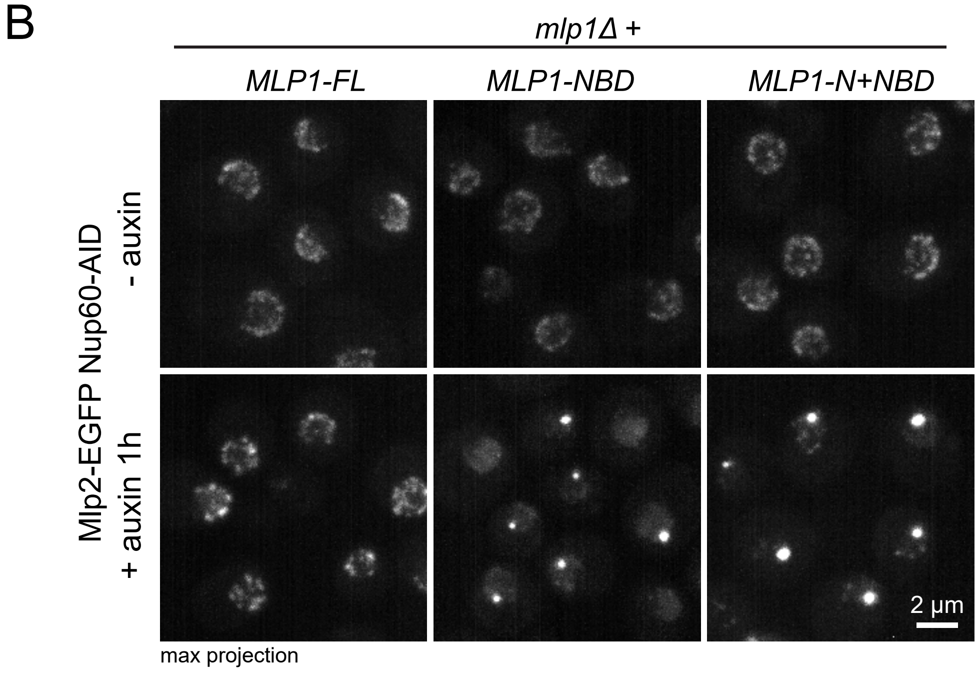


**Figure S3: Mlp1 and Mlp2 stabilize each other at the NPC in the absence of Nup60.**
(A) Localization of Mlp1-mNG upon rescue of *mlp2Δ* with different Mlp2 truncation constructs. Cells were treated with auxin for 1h to deplete Nup60.
(B) Localization of Mlp2-mNG upon rescue of with different Mlp1 truncation constructs. Cells were treated with auxin for 1h to deplete Nup60.

### Figure S4

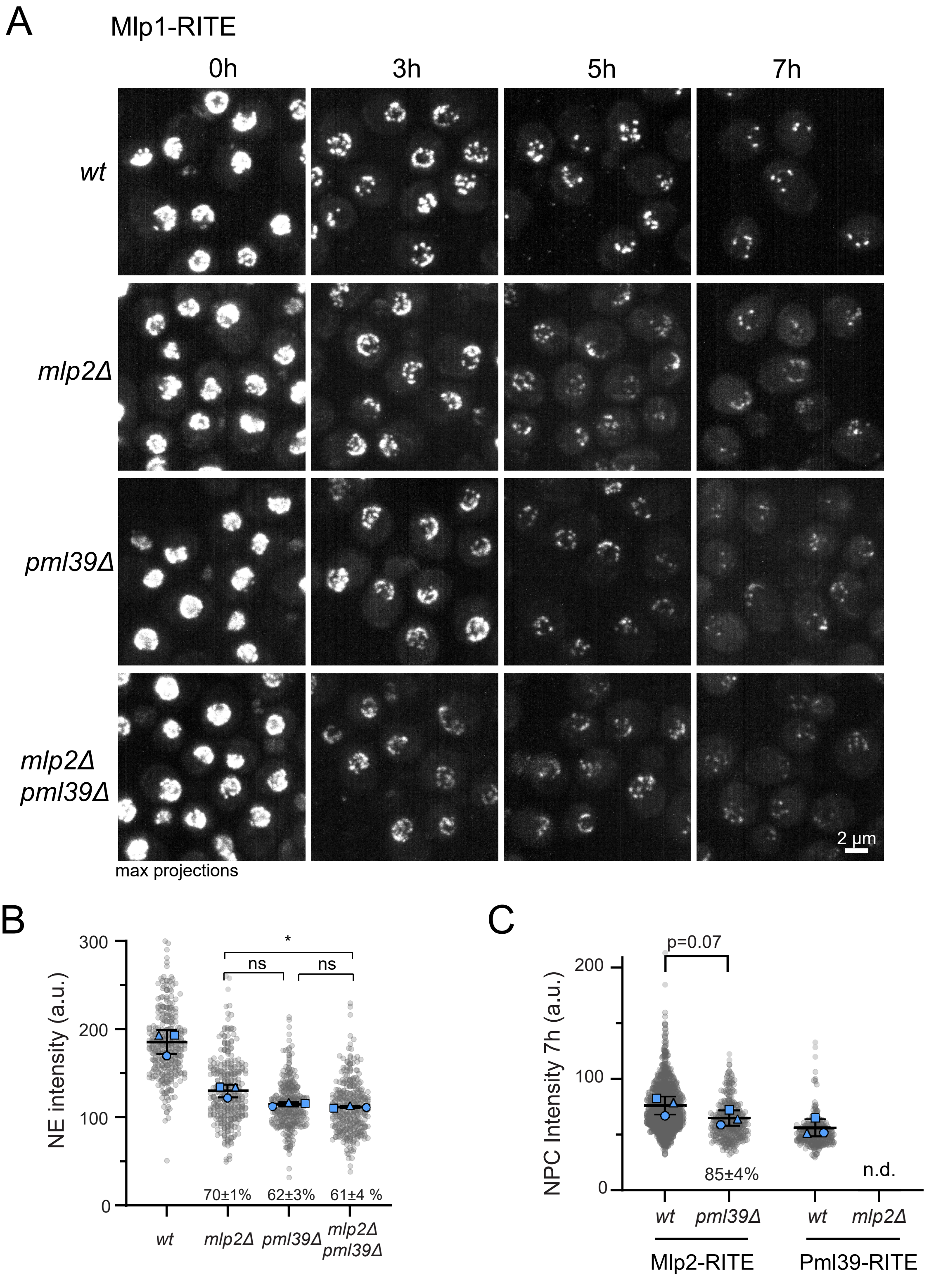


**Figure S4: Mlp2 contributes to anchoring of Mlp1 to the NPC.**

(A) Additional timepoints for the RITE time course shown in Figure 4A. Maximum intensity projections are shown.

(B) Quantification of NE intensity at 0h timepoint for data shown in Figure 4A. Grey circles represent background-subtracted measurements from individual cells. Blue symbols represent means of individual biological replicates. Error bars represent the standard deviation. Statistical analysis was carried out a repeated measures one-way ANOVA with Tukey's multiple comparison test. ns: not significant, * $p < 0.05$. All other comparison (wt with each mutant) are significant with $p \leq 0.0001$. Percentage values indicate the relative mean intensity compared to wt ± standard deviation.

(C) Quantification of mean NPC intensity at the 7h timepoint for data shown in Figure 4C and 4D. Grey circles represent background-subtracted measurements from >75 individual NPCs per biological replicate. Blue symbols represent means of individual biological replicates. Error bars represent the standard deviation. p value is for a two-tailed paired t-test. Percentage value indicates the relative mean intensity compared to wt ± standard deviation.

### Figure S5

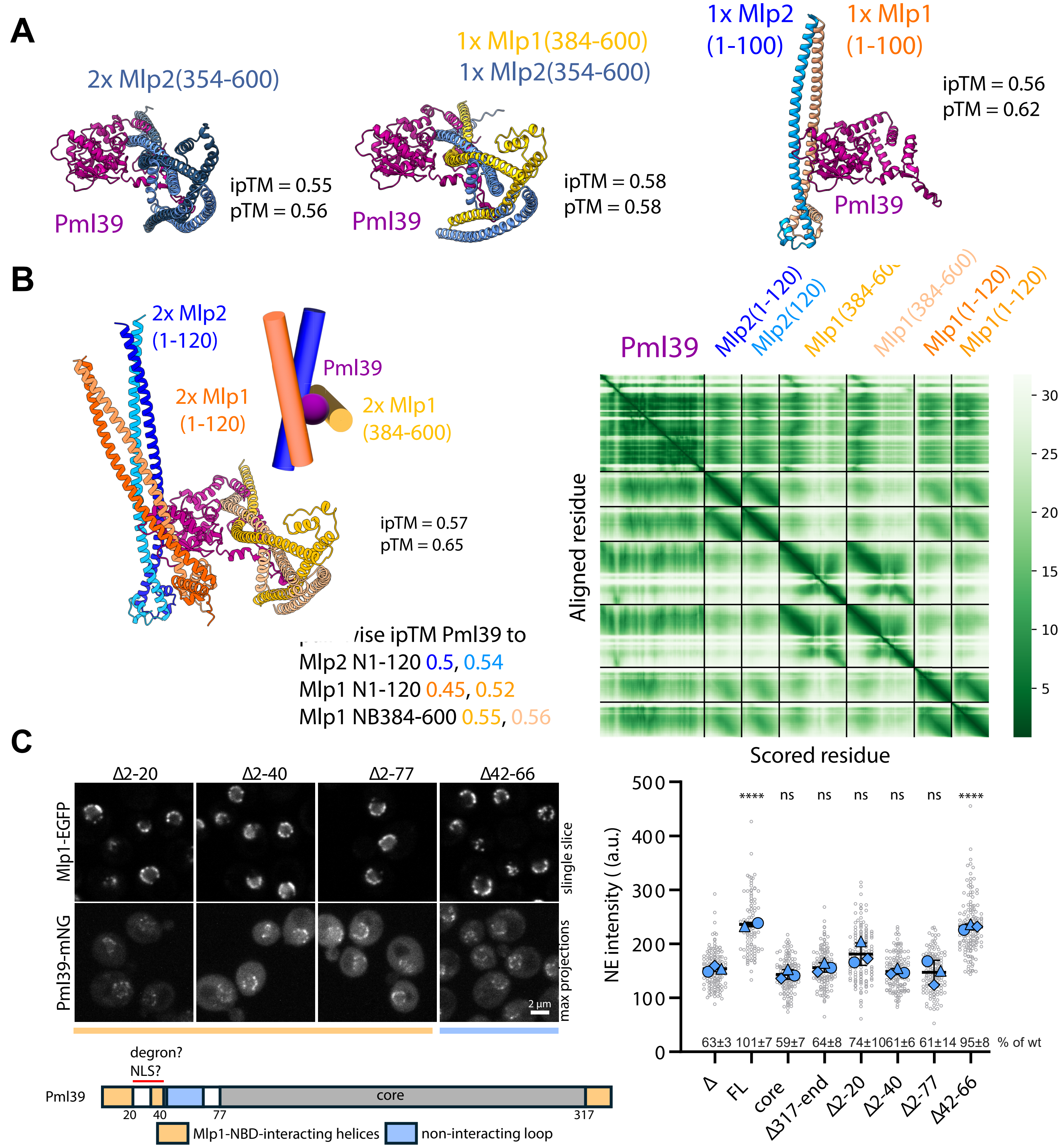


**Figure S5: Investigating the interaction network of Pml39 at the NPC.**
(A) AlphaFold3 model of the interaction of Pml39 with the NPC binding domain of Mlp1 and the N-termini of Mlp1 and Mlp2. Overall AlphaFold scores are given.
(B) AlphaFold3 model of the interaction of Pml39 with the NPC binding domain of Mlp1 and the N-termini of Mlp1 and Mlp2. Schematic on top right illustrates orientation of proteins toward each other. The matrices of the PAE scores is shown on the right. The overall ipTM and pTM scores as well as the pairwise ipTM scores are shown below.
(C) Localization of Mlp1-EGFP in strains expressing additional Pml39 truncation mutants. Plot shows quantification of Mlp1-EGFP intensity at the nuclear envelope. Grey circles represent background-subtracted measurements from individual cells (~50 cells per condition and biological replicate). Blue symbols represent means of individual biological replicates. Error bars represent the standard deviation. Statistical testing was performed by fitting a mixed effects model with Sidak's multiple comparison test comparing each strain to Δ. **** < 0.0001 , ns > 0.05. Percentage values indicate the relative mean intensity compared to FL ± standard deviation. Note that the quantification data for the strains shown in Fig. 5E is shown again for comparison. Schematic indicates the different deleted regions on the Pml39 architecture. A region that appears to affect the expression level of the truncation mutants is indicated in red.

### Figure S6

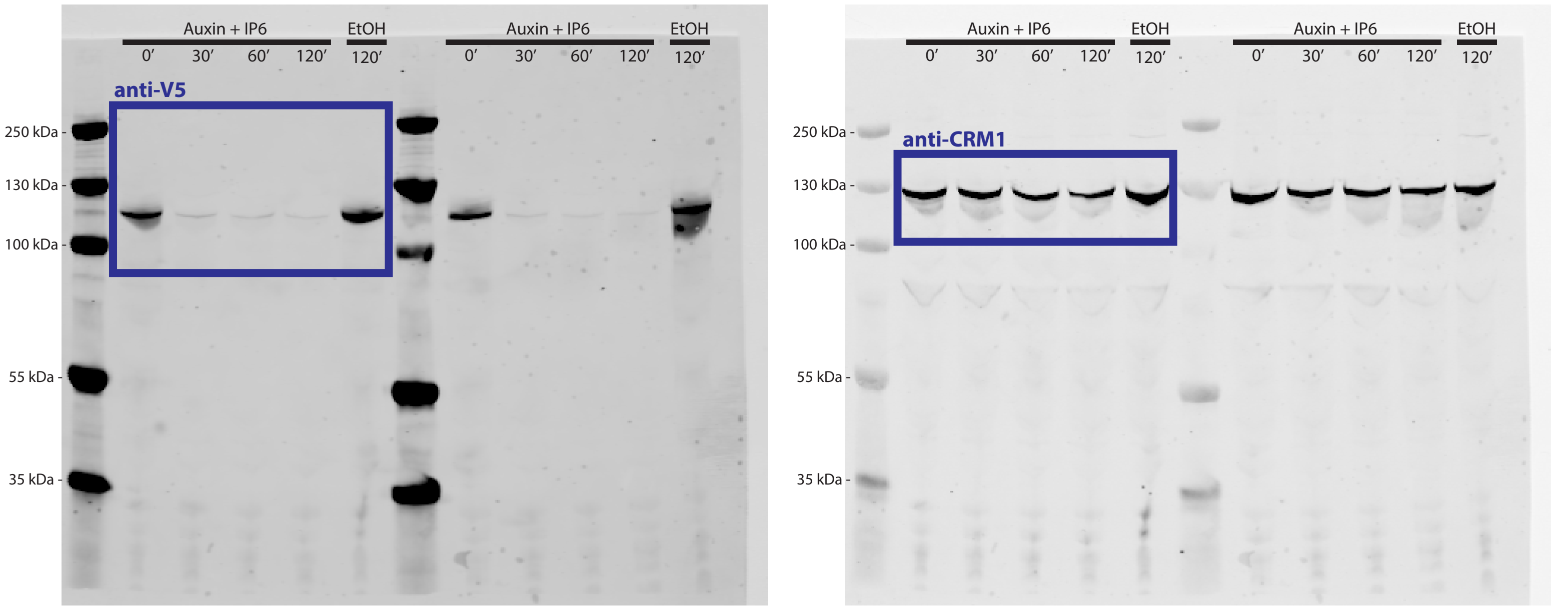


**Figure S6: Blot Transparency.**
Full Western blots of blots used in Fig.S2A are shown.
